# Telomeric repeat evolution in the phylum Nematoda revealed by high-quality genome assemblies and subtelomere structures

**DOI:** 10.1101/2023.05.24.542215

**Authors:** Jiseon Lim, Wonjoo Kim, Jun Kim, Junho Lee

**Affiliations:** Department of Biological Sciences, Seoul National University, Gwanak-ro 1, Gwanak-gu, Seoul 08826, Korea; Institute of Molecular Biology and Genetics, Seoul National University, Seoul 08826, Korea; Research Institute of Basic Sciences, Seoul National University, Seoul 08826, Korea; Department of Convergent Bioscience and Informatics, College of Bioscience and Biotechnology, Chungnam National University, Daehak-ro 99, Daejeon 34134, Republic of Korea

## Abstract

Telomeres are composed of tandem arrays of telomeric-repeat motifs (TRMs) and telomere-binding proteins (TBPs), which are responsible for ensuring end-protection and end-replication of chromosomes. TRMs are highly conserved due to the sequence specificity of TBPs, but significant alterations in TRM have also been observed in several taxa, except Nematoda. We used public whole-genome sequencing datasets to analyze putative TRMs of 100 nematode species and determined that two distinct branches included specific novel TRMs, suggesting that evolutionary alterations in TRMs occurred in Nematoda. We focused on one of the two branches, the Panagrolaimidae family, and performed a *de novo* assembly of four high-quality draft genomes of the canonical (TTAGGC) and novel TRM (TTAG<u>A</u>C)-containing isolates; the latter genomes revealed densely clustered arrays of the novel TRM. We then comprehensively analyzed the subtelomeric regions of the genomes to infer how the novel TRM evolved. We identified DNA damage–repair signatures in subtelomeric sequences that were representative of consequences of telomere maintenance mechanisms by alternative lengthening of telomeres. We propose a hypothetical scenario in which TTAG<u>A</u>C-containing units are clustered in subtelomeric regions and pre-existing TBPs capable of binding both canonical and novel TRMs aided the evolution of the novel TRM in the Panagrolaimidae family.

## INTRODUCTION

Since linear chromosomes originated from a circular structure, a specific DNA–protein complex has evolved to protect the ends of the chromosomes, i.e. the telomere. Telomeres are typically composed of tandem arrays of telomeric-repeat motifs (TRMs) and telomere-binding proteins (TBPs) that can bind to the tandem arrays (Zhong et al. 1992). TBP binding interferes with the accession of other proteins. Hence, proteins that respond to DNA damage signals cannot bind to the exposed ends of linear chromosomes and the exposed ends are not recognized as DNA damage sites (Van Steensel et al. 1998; De Lange 2009; Lazzerini-Denchi and Sfeir 2016). This enables the telomere to maintain chromosome integrity.

TRM sequences are typically well conserved because TBPs attach to arrays of TRMs in a sequence-specific manner. However, dozens of variant TRMs have evolved in many taxa, including ≥150 motifs in fungi (Červenák et al. 2021) and ≥7 motifs in plants (Peska and Garcia 2020). Furthermore, except Hymenoptera, many arthropod species have unique telomere structures. For example, most Insecta species have telomere structures composed of telomere-specific retrotransposons interposed between canonical TRMs, whereas Diptera species have telomeres that are exclusively composed of retrotransposons and lack TRMs (Abad et al. 2004; Garavís et al. 2013; Zhou et al. 2022). It is yet to be elucidated how TRMs have been altered while interacting with TBPs. These diverse TRMs and telomere structures developed from the co-evolution of a TRM and several TBPs (Shakirov et al. 2009; Steinberg-Neifach and Lue 2015; Sepsiova et al. 2016; Červenák et al. 2019); however, they may have also originated from TRM-specific changes regardless of TBP changes if TBPs were capable of binding to several types of TRMs (Rotková et al. 2004; Fajkus et al. 2005; Kramara et al. 2010; Visacka et al. 2012; Tomáška et al. 2019; Červenák et al. 2021).

We focused our analysis on subtelomeric regions and the DNA damage–repair signatures generated during telomere evolution, especially telomere and subtelomere reconstruction by alternative lengthening of telomeres (ALT) mechanisms (Heaphy et al. 2011; Sobinoff and Pickett 2017). ALT refers to alternative mechanisms that substitute telomerase function for telomere maintenance at the chromosome ends (Bryan et al. 1995). It becomes remarkably functional in telomerase mutant eukaryotic cells such as those in yeast, *Caenorhabditis elegans*, mouse, and humans, and can thus be used to model and interpret the evolution of TRM changes (Lundblad and Blackburn 1993; Bryan et al. 1995; Seo et al. 2015; Kim et al. 2020; Kim et al. 2021a; Kim et al. 2021b). In these ALT models, homology-directed repair (HDR) mechanisms can add new telomeric components. HDRs exploit the homology between shortened telomeric sequences and tandem arrays of TRMs in other genomic loci to replicate the tandem array and adjacent sequences to the shorter end (Lydeard et al. 2007; Roumelioti et al. 2016; Kramara et al. 2018; Kim et al. 2020). Furthermore, a unique sequence inserted between TRMs, known as the template for ALT, may be used to repair shorter telomeric repeats even in wild-type *C. elegans*, resulting in the formation of a new subtelomeric structure (Kim et al. 2019a). These findings suggest that ALT mechanisms play a role in the evolution of telomeres and the evolutionary route can be reconstructed by studying traces left behind ALT activities in subtelomeric sequences.

The present study explored TRM evolution in Nematoda using public whole-genome sequencing (WGS) data obtained from 100 nematode species. We determined that TTAGGC, a canonical Nematoda TRM, had changed in two branches and validated that TTAGAC had evolved in the family Panagrolaimidae by sampling 14 isolates and conducting WGS, followed by the generation of high-quality genome assemblies. We evaluated the subtelomeric regions of TTAGGC-telomere isolates of Panagrolaimidae to infer possible evolutionary paths toward TTAGAC telomeres.

## RESULTS

### Identification of two novel TRMs in Nematoda

We examined variations in TRMs using public short-read WGS data from Nematoda species that have matched public genome assemblies from all five major clades. A total of 100 species met the criteria, including at least 17 species from Clades I, III, IV and V but only one from Clade II (Fig. 1A). The species and accession numbers used in this analysis are listed in Supplemental Table S1 and S2.

**Figure 1.**
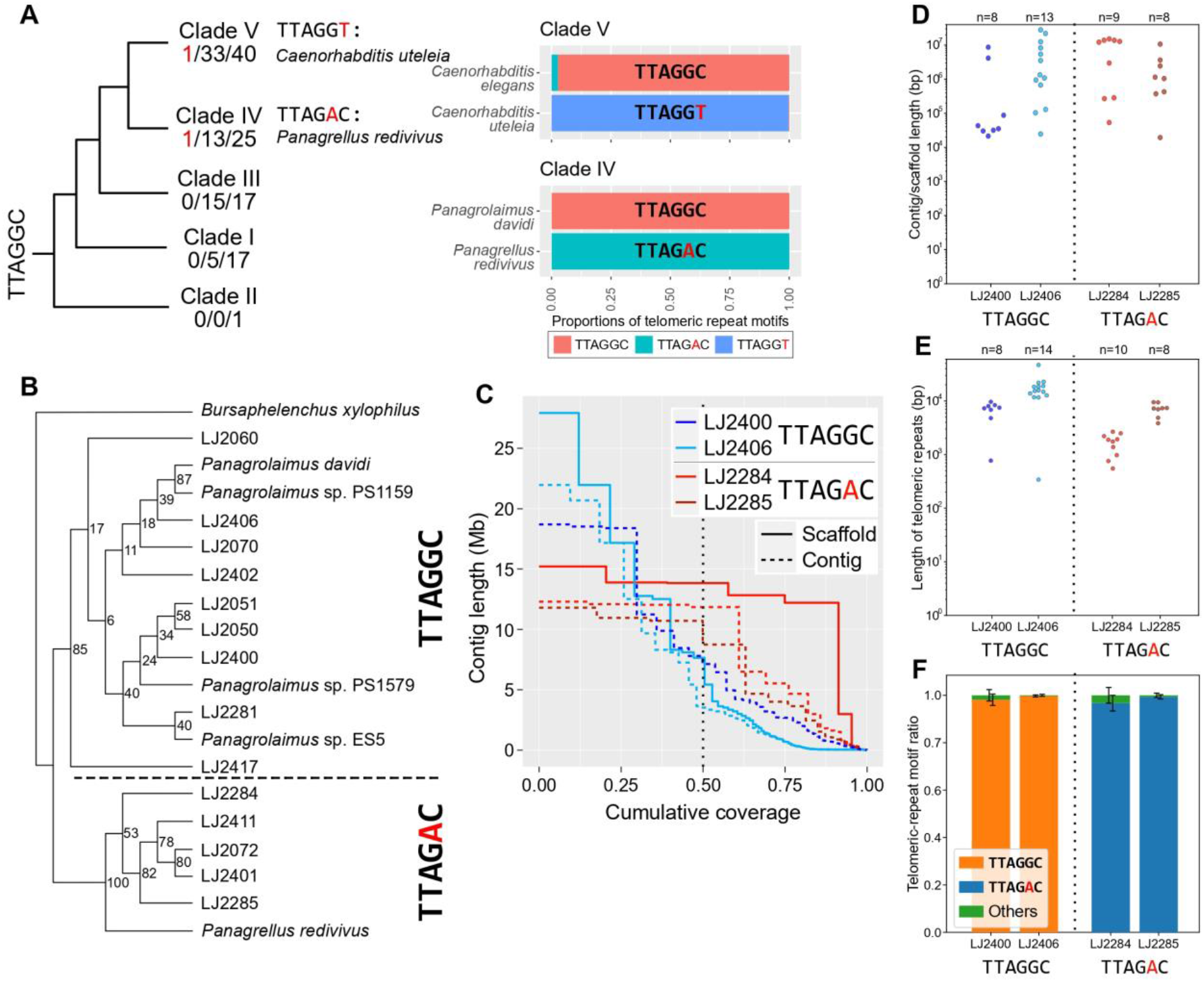
Putative novel TRMs in the phylum Nematoda and validation of the emergence of TTAGAC in the family Panagrolaimidae. (A) The cladogram was adapted from Smythe et al. (Smythe et al. 2019). The three numbers separated by ‘/’ under each clade indicates the number of species with novel TRMs/the number of species whose TRM was identified in our analysis/total number of species we analyzed, respectively. Changes from TTAGGC, the canonical Nematoda TRM, to the novel TRMs are highlighted in red. Below each novel TRM is a species name with the corresponding TRM. Each bar indicates the proportion of the canonical Nematoda TRM (TTAGGC) and novel TRMs in each species. The upper bar of each pair represents a control species that harbors the canonical TRM. The TRMs are represented as follows: red, TTAGGC; cyan, TTAGAC; blue, TTAGGT. (B) Phylogenetic relationships of 19 Panagrolaimidae species/isolates based on their 18S rDNA sequences and their putative TRMs. *Bursaphelenchus xylophilus* was used as an outgroup species. The number on each node represents the bootstrap support value. (C) Contig/scaffold length distributions of the genome assemblies. The vertical dotted line indicates N50 contig/scaffold size. (D–F) The vertical dotted line separates the TTAGGC-telomere isolates (left) and the TTAGAC-telomere isolates (right). (D) Length distributions of the contigs/scaffolds containing highly clustered telomeric repeats at the end. Each dot represents each contig/scaffold, and the total number of contigs/scaffolds of each isolate (n) is indicated at the top of the graph. (E) Estimated length distributions of clustered telomeric repeats at the end of the contigs/scaffolds. Each dot indicates a telomeric-repeat cluster for each contig/scaffold, and the total number of telomeric-repeat clusters for each isolate (n) is indicated at the top of the graph. (F) The proportion of TRM types in clustered telomeric repeats. Error bars represent the standard deviation for all clustered telomeric repeats at the end of contigs/scaffolds in each isolate.

The canonical TRM of Nematoda is known to be a 6-bp sequence (TTAGGC) that covers a >1-kb length in *C*. *elegans* telomeres (Kim et al. 2019a; Yoshimura et al. 2019). We examined the WGS data from 100 species to identify novel TRMs with a concatemer of 5–7 bp, a high copy number and a pattern similar to TTAGGC (Supplemental Table S1 and S2). We were unable to determine the TRM in 34 species because of their low copy number motifs, very short TRM array or contamination of sequenced samples (Supplemental Table S2 and Supplemental Note). TTAGGC, the canonical Nematoda TRM, was identified in 64 of the remaining 66 species (Fig. 1A and Supplemental Table S1).

Interestingly, we determined that the other two nematode species had few TTAGGC copies but a high copy number of 1 of 2 putative TRMs: TTAGGT or TTAGAC (Fig. 1A, Supplemental Fig. S1 and Supplemental Tables S1 and S3; see Supplemental Note and Supplemental Table S4 for further validation of the TRMs). The identified TRMs were new to Nematoda and had not been reported previously. We assumed that these two motifs evolved independently because they occurred in different clades.

### The emergence of a new TRM, TTAGAC, in the family Panagrolaimidae

We focused on the TTAGAC sequence of the family Panagrolaimidae to confirm the noncanonical TRM sequence identified by short-read WGS data because many different isolates of this family were widely isolated from regions across the Republic of Korea. Some were easily cultured under laboratory conditions. We used 19 short-read WGS datasets of the family Panagrolaimidae to determine whether the unique motif TTAGAC emerged once or multiple times (Supplemental Table S5). WGS datasets of five species were available from publicly accessible databases, whereas the remainder were derived from sequencing data of the isolates collected from the Republic of Korea, which were probably distinct species as they had distinguishable ribosomal DNA (rDNA) sequences (Supplemental Table S6). *B. xylophilus* was also used as the outgroup species.

We determined that *Panagrellus redivivus* and five out of fourteen collected isolates had the novel TRM, TTAGAC, whereas all four *Panagrolaimus* species and the remaining nine isolates possessed the canonical TRM, TTAGGC. Moreover, the five collected TTAGAC-TRM isolates clustered with *Panagrellus redivivus*; however, the nine TTAGGC-TRM isolates were grouped with the *Panagrolaimus* species (Fig. 1B and Supplemental Table S5). Furthermore, the TTAGGC concatemer was not observed in any of the 6 TTAGAC-TRM species/isolates, nor was the TTAGAC concatemer found in any of the 13 TTAGGC-TRM species/isolates. These results suggest that TTAGAC sequences in Panagrolaimidae may have arisen from a single ancestor that had TTAGAC as its TRM.

### High-quality genome assemblies confirmed that TTAGAC repeats were clustered at the ends of contigs or pseudo-chromosome molecules in TTAGAC-TRM isolates

We constructed *de novo* genome assemblies of Panagrolaimidae isolates using PacBio high-fidelity (HiFi) long-read sequencing technology and Arima Hi-C technology to assess whether the newly identified TRM presented clusters at the ends of the chromosomes. Two TTAGAC-TRM isolates, LJ2284 and LJ2285, and two TTAGGC-TRM isolates, LJ2400 and LJ2406, were selected for the study. We produced 3.1 Gb (43×), 2.9 Gb (55×), 17.4 Gb (26×) and 36.1 Gb (93×) HiFi reads for LJ2284, LJ2285, LJ2400 and LJ2406, respectively. The HiFi reads were then assembled into high-quality draft genomes in which the length of the longest contig and contig N50 varied from 11.80 Mb to 21.96 Mb and 3.28 Mb to 11.84 Mb, respectively. We also generated Hi-C data for LJ2284 and LJ2406 and scaffolded their contigs into larger chunks (Supplemental Fig. S2 and Supplemental Table S7). The longest scaffold lengths increased to 15.2 Mb and 27.9 Mb, and their scaffold N50 increased to 13.8 Mb and 7.6 Mb for LJ2284 and LJ2406, respectively (Fig. 1C and Supplemental Table S8). Additionally, 91.4% of the LJ2284 contigs were connected as five scaffolds (Supplemental Fig. S2).

All of our genome assemblies had ∼75% BUSCO completeness, which was comparable to that of the nearly complete genome assembly of *B*. *xylophilus* (71%) (Supplemental Fig. S3) (Dayi et al. 2020). In addition, we analyzed the synteny relationships between *B*. *xylophilus* and our genome assemblies using their common single-copy orthologs. Pseudo-chromosome-level scaffolds of LJ2284 and contigs of LJ2285 exhibited marked collinearity for all the chromosomes, except chromosome 4 of *B*. *xylophilus* (Supplemental Fig. S4 and Supplemental Note). Chromosomes 5 and 6 of *B*. *xylophilus* also showed collinearity with clusters of LJ2400 contigs and LJ2406 scaffolds, but the collinearity of other chromosomes was much weaker (Supplemental Fig. S4 and Supplemental Note). Based on these findings, our Panagrolaimidae genome assemblies were highly contiguous.

The constructed high-quality genome assemblies revealed that the telomeric repeats of both LJ2284 and LJ2285 changed from TTAGGC to TTAGAC. First, we confirmed whether the telomeric repeats had been adequately assembled. Our analysis revealed that each genome assembly had 8–13 contigs/scaffolds ending with telomeric repeats (Fig. 1D). Among these contigs/scaffolds, one each of LJ2284 and LJ2406 was assembled at the telomere-to-telomere level (Supplemental Tables S7 and S8). The telomeric repeats at the ends of the contigs/scaffolds in each genome assembly exhibited various ranges of length in the Panagrolaimidae isolates (Fig. 1E and Supplemental Table S9). Their lengths in LJ2284 ranged from 0.5 to 2.7 kb, but they were much longer in LJ2285 and LJ2400, ranging from 3.8 to 9.7 kb (except for one in LJ2400). LJ2406 exhibited telomeric-repeat clusters longer than 10 kb (except for one cluster). Because 84.9%–99.4% of raw HiFi reads are longer than 10 kb, the TRM cluster lengths at each contig/scaffold end would be highly correlated with their actual lengths (Supplemental Fig. S5). The lengths of the clustered TRMs were comparable to those of the two *C*. *elegans* assemblies constructed using long-read sequencing technologies (2.3– 5.7 kb) (Kim et al. 2019a; Yoshimura et al. 2019). Furthermore, the telomeric repeats consisted mainly of TTAGAC in LJ2284 (96.67%) and LJ2285 (99.35%) and of TTAGGC in LJ2400 (98.11%) and LJ2406 (99.73%) (Fig. 1F, Supplemental Fig. S6 and Supplemental Table S9). These findings supported our TRM study employing short-read WGS data and revealed that the TTAGAC telomere had evolved in the family Panagrolaimidae.

### TTAGAC-containing units clustered near telomeric regions of TTAGGC-telomere isolates

Because the most nematode species that we analyzed had a canonical Nematoda TRM sequence, TTAGGC, the TTAGAC telomere probably evolved from its ancestral TTAGGC telomere via a single nucleotide mutation of the telomerase RNA component gene, *terc*. We hypothesized that subtelomeric sequences can provide evidence regarding the evolutionary mechanism because these can carry historical records of telomere dynamics. For example, it has previously been documented that ALT mechanisms have been used to reconstruct subtelomeric and telomeric regions in *C. elegans* and that templates for ALT in subtelomeric regions can be used to repair telomere damage or maintain the integrity of the telomere (Seo et al. 2015; Kim et al. 2019a; Kim et al. 2020; Kim et al. 2021b; Lee et al. 2022b). Thus, we analyzed subtelomeric regions of our four Panagrolaimidae genome assemblies to trace ancestral telomere damage and repair events that could explain how the TTAGAC telomere evolved from the TTAGGC telomere.

Interestingly, most telomeric repeats in telomeric regions (7/10 in LJ2284, 4/8 in LJ2285, 6/8 in LJ2400 and 8/14 in LJ2406) were attached to unit clusters containing TTAGGC or TTAGAC (Fig. 2A and Supplemental Table S10). This finding was striking, as none of the subtelomeric regions of *C*. *elegans* and *B*. *xylophilus* exhibited TRM-containing unit clusters directly attached to the telomeric regions (Fig. 2A and Supplemental Table S11). Furthermore, both TTAGGC-containing unit clusters and TTAGAC-containing unit clusters were identified in all genomes of the TTAGGC- and TTAGAC-telomere isolates, with the exception of the LJ2285 genome (TTAGAC telomere), which contained only TTAGAC-containing unit clusters. For LJ2284, LJ2400 and LJ2406, these clusters consisted of repetitive units with length distributions ranging from 14 bp to ∼2 kb, and these repetitive units were tandemly repeated on an average of ≥19 copies (Supplemental Tables S10 and S12). The LJ2285 genome did not contain such same-length unit clusters; however, it contained sequence clusters consisting of 5–9 copies of units containing TTAGAC and similar sequences (10–74 bp). These TRM-containing unit clusters were not found outside the subtelomeric region in the telomere-containing contigs/scaffolds (see also Supplemental Note describing a contrasting case in *C*. *elegans*), raising the possibility that these clusters are associated with telomere maintenance in Panagrolaimidae. Notably, we also identified a similar subtelomeric pattern in *Caenorhabditis uteleia*, the noncanonical TTAGGT-TRM species, where TTAGGT- and/or TTAGGC-containing unit clusters were attached to or close to long TTAGGT repeat arrays (Supplemental Fig. S7 and Supplemental Note).

**Figure 2.**
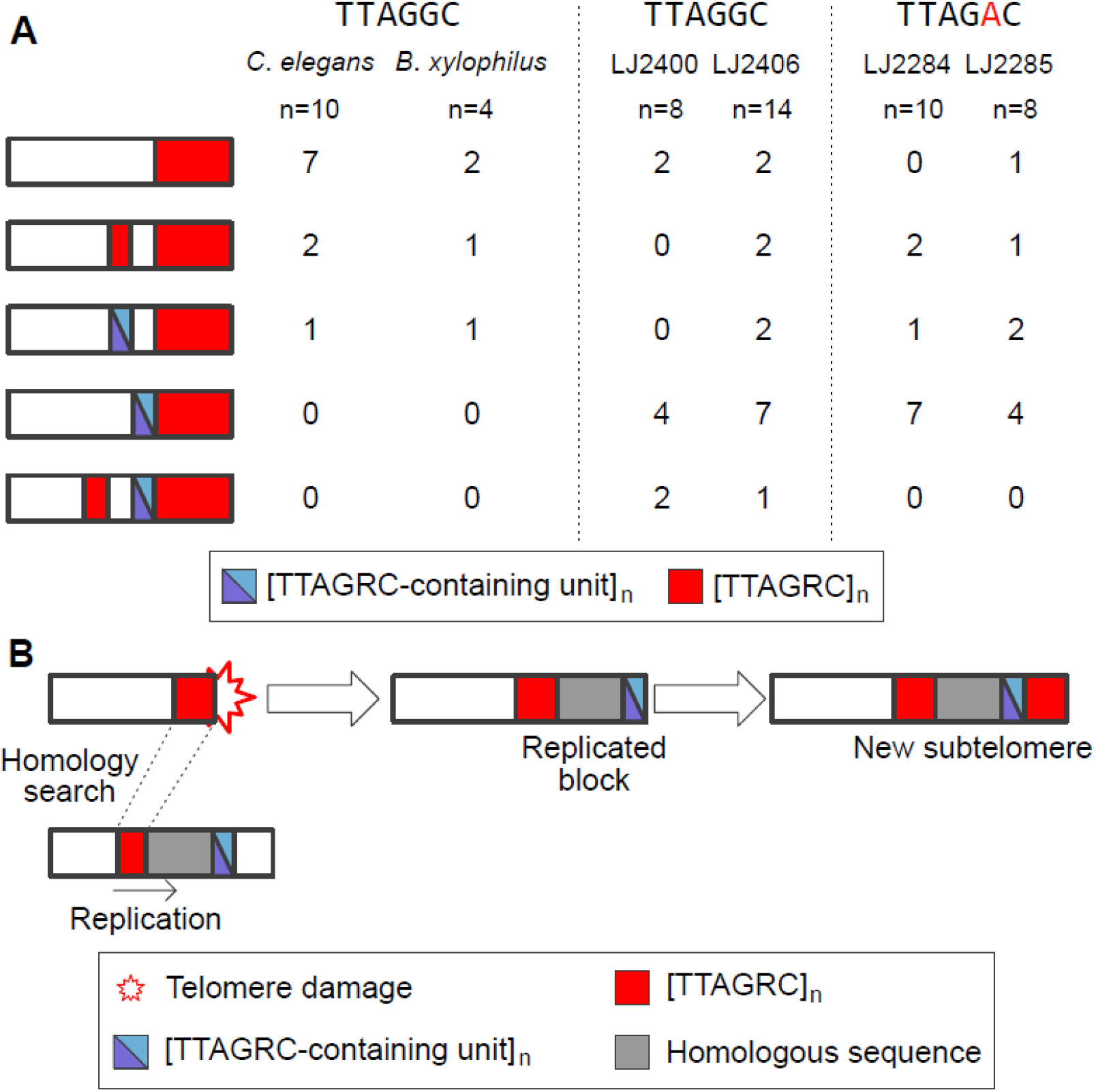
Subtelomere structures and a proposed model to explain clustered blocks of TRMs or TTAGRC-containing unit clusters in the subtelomeric regions. (A) Schematic representation of subtelomere structures in the contigs/scaffolds listed in Supplemental Table S10. We categorized the subtelomere structure into a total of five types based on (i) whether it has ITS (shorter red blocks) and/or TTAGRC-containing unit clusters (half-sky blue, half-violet blocks) and (ii) whether ITS, unit clusters and telomeric regions were directly attached or separated by other sequences (white, empty blocks). Each value refers to the number of contigs/scaffolds with the corresponding structure in each species/isolate. A detailed classification of subtelomere structures into 12 subtypes is shown in Supplemental Fig. S8. (B) A proposed model for generating subtelomere structures containing ITS or unit clusters. After telomere damage, HDR mechanisms exploit homology between shortened telomeric repeats and clusters of variant telomeric repeats, including TTAGRC-containing unit clusters, to repair the damaged telomere. A HDR mechanism, BIR, may replicate sequences near the cluster with variant telomeric repeats, creating a new homologous block of the original sequence block. If the original template block exhibits a unit cluster, the cluster would also be replicated.

In addition to the TRM-containing unit clusters, interstitial telomeric sequences (ITS) were frequently observed in the subtelomeric regions of the constructed genome assemblies, which were consistent with the ITS enrichment present in *C*. *elegans* chromosome arms (Consortium* CeS 1998). All subtelomere structures were categorized into five different subtelomere structures based on the location of the unit clusters and/or the ITS (Fig. 2A and Supplemental Table S10; see also Supplemental Fig. S8). The four genome assemblies exhibited similar patterns. For most of the subtelomere structures in the four genome assemblies, the TRM-containing unit cluster was positioned directly adjacent to the telomeric region. The simplest subtelomere structure, which did not contain any unit cluster or ITS, accounted for fewer than a quarter of all the subtelomere structures. In contrast, the remaining subtelomere structures had at least one unit cluster or ITS separated from the telomeric region, which may have resulted from ancestral telomere damage and repair processes (Fig. 2B).

### TTAGAC-containing units were probably used to maintain the integrity of TTAGGC telomeres

We postulated that the identified subtelomeric clusters of the TTAGRC-containing units and ITS were evidence of ancestral telomere damage and repair via ALT mechanisms and that if true, this would help understand the role of TTAGRC-containing unit clusters in telomere maintenance (Fig. 2B). We focused on BIR, one of the most well-known ALT mechanisms for telomere repair (Malkova et al. 1996; Bosco and Haber 1998; Kim et al. 2019a; Kim et al. 2020). Telomeres that have been damaged or shortened should be repaired, and BIR may rely on homology between shorter telomeric repeats and variant telomeric repeats at other loci (Fig. 2B, left side). The replication was then activated, and occasionally, sequences beside the variant telomeric repeats would also be replicated (Fig. 2B, middle). Furthermore, if the original homology block contained a TTAGRC-containing unit cluster, the cluster would also be duplicated simultaneously, resulting in a new subtelomere structure with a homology block interposed between the ITS and the TTAGRC-containing unit cluster (Fig. 2B, right side). We then hypothesized that if the subtelomere structure (ITS, a sequence block, and a TTAGRC-containing unit cluster) of our genomes had been generated via BIR, we should be able to identify another sequence block homologous to the interposed sequence between the ITS and the TTAGRC-containing unit cluster.

To test this hypothesis, we first identified subtelomeres with a TTAGRC-containing unit cluster located between ITS and a telomeric region. A total of three contigs were identified: ptg000074l and ptg000145l of LJ2400 and ptg000247l of LJ2406 (Fig. 3). Interestingly, ptg000074l and ptg000145l of LJ2400 shared 8-kb and 9-kb homology blocks adjacent to their ITS and had a TTAGGC-containing unit cluster ∼9-kb and ∼200-bp away from the shared block, respectively. These 8-kb and 9-kb homology blocks pointed to the possibility of their replication via BIR using the ITS as homology.

**Figure 3.**
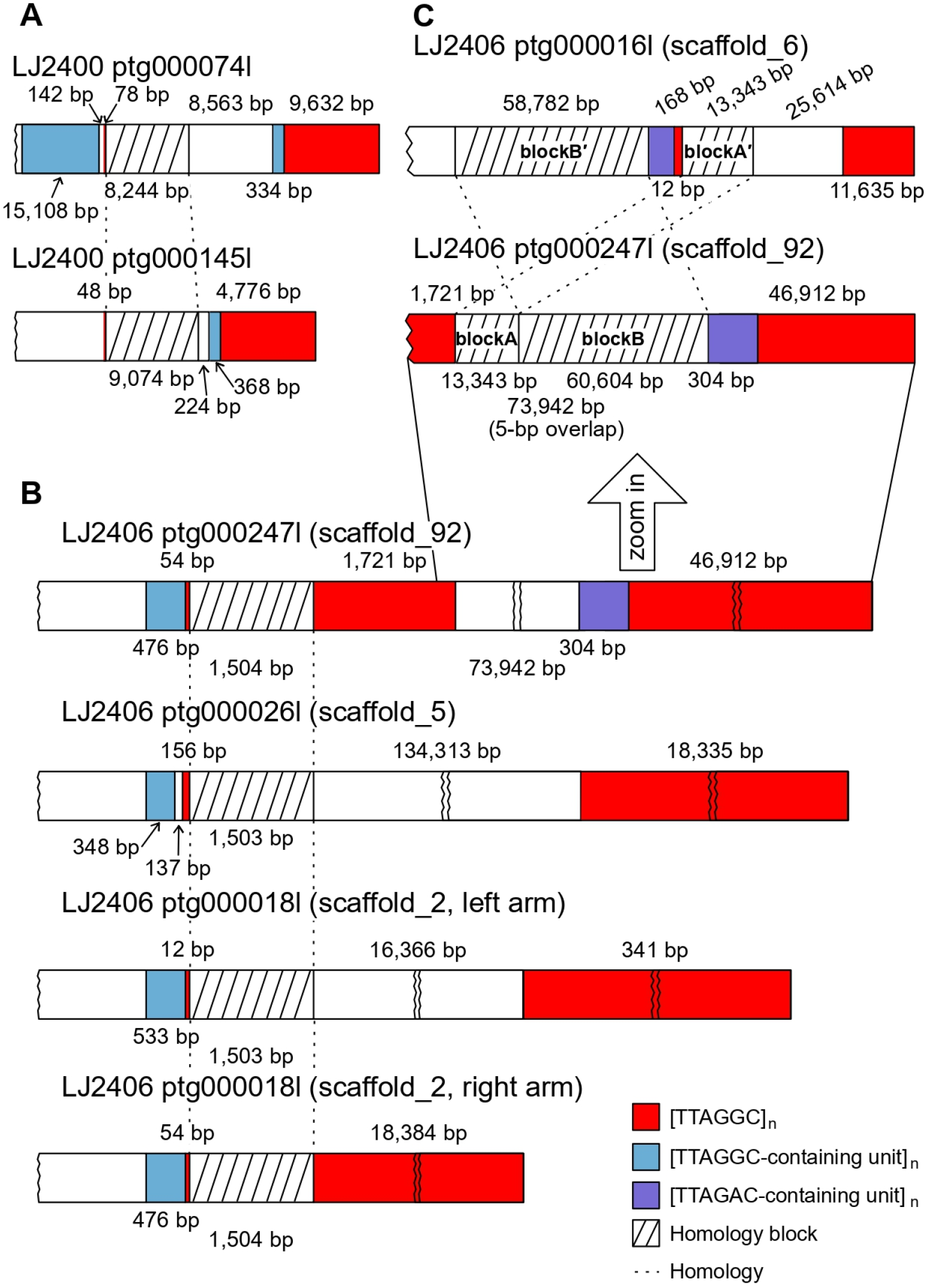
Traces of telomere damage and repair processes through ALT mechanisms identified in subtelomeric regions of genome assemblies of TTAGGC-telomere isolates. (A–C) Schematic representations of subtelomeres in the genome assemblies of LJ2400 and LJ2406. Hatched boxes indicate homology blocks between two or more subtelomeres, and dashed lines indicate homologous relationships. The red boxes represent telomeric-repeat clusters. Violet and sky blue boxes represent TTAGAC-containing unit clusters and TTAGGC-containing unit clusters, respectively. Not to scale.

The ptg000247l of LJ2406 had a more evident BIR signature than the ptg000074l and ptg000145l of LJ2400, indicating that the subtelomeric TTAGRC-containing unit cluster was built via the ALT process. The subtelomeric region of ptg000247l of LJ2406 consisted of a TTAGGC-containing unit cluster, a 54-bp ITS, a 1.5-kb sequence block, a 1.7-kb ITS, a 74-kb sequence block and a TTAGAC-containing unit cluster close to the telomeric region (Fig. 3B). Interestingly, by searching for this complex subtelomeric sequence in the whole genome using BLAST, we determined that the 1.5-kb and 74-kb blocks had homologous sequences in other subtelomeric regions. The 1.5-kb block between the 54-bp ITS and 1.7-kb ITS had almost identical homologous sequences in three different subtelomeric regions. These shared blocks were directly adjacent to short and variable-length ITS near TTAGGC-containing unit clusters (Fig. 3B). The TTAGGC-containing cluster and ITS may have been utilized for homology to replicate the 1.5-kb homology blocks.

The 74-kb block between the 1.7-kb ITS and the TTAGAC-containing unit cluster possessed another trace of HDR and an additional replication of a TTAGRC-containing unit cluster after telomere damage (Fig. 3C). As the 1.7-kb length of the ITS is too long to be created by random mutation, it could represent evidence of ancestral telomere damage and repair. The 74-kb block contained at least two distinct homology blocks: the 13-kb blockA and the 61-kb blockB (Fig. 3C). The 13-kb blockA directly linked to the ITS in ptg000247l had an identical blockA′ in ptg000016l that was also directly attached to another 180-bp sequence of ITS and a TTAGAC-containing unit cluster. The 61-kb blockB was homologous but not identical to the 59-kb blockB′ located on the opposite side of the 180-bp block of the ITS and the TTAGAC-containing unit cluster in ptg000016l. Of note, the last 5 bp of the 13-kb blockA and the first 5 bp of the 61-kb blockB were identical (i.e. TAAAT). The 61-kb blockB was further divided into 4 blocks, each of which also exhibited microhomologies with the adjacent blocks (Supplemental Fig. S9 and Supplemental Table S13). These data imply that telomere damage and repair may have occurred in the following order: first, the 13-kb blockA in ptg000247l was replicated by BIR using homology between the 1.7-kb ITS in ptg000247l and the 180-bp block of ITS and a TTAGAC-containing unit cluster in ptg000016l. Next, the 61-kb blockB in ptg000247l was replicated by multiple rounds of microhomology-mediated BIR (MMBIR) and template switching events using microhomology sequences at the end of the blocks (Supplemental Fig. S9 and Supplemental Note).

This 59-kb blockB′ in ptg000016l was directly attached to the TTAGAC-containing unit cluster, which was duplicated in ptg000247l. Notably, the original TTAGAC-containing unit cluster in ptg000016l was shorter than the duplicated cluster in ptg000247l, implying that the duplicated cluster nearly doubled after MMBIR was completed. The duplicated TTAGAC-containing unit cluster in ptg000247l was directly linked to the TTAGGC telomere, indicating that the duplicated cluster was exposed at the end. These lines of evidence support the notion that even in TTAGGC-telomere species, TTAGAC-containing units could be incorporated to constitute a telomere or at least serve as a DNA template for replenishing TRM repeats by telomerase. This use of TTAGAC-containing units in the TTAGGC-telomere isolate may have contributed to the evolution of the TTAGAC telomere from the TTAGGC telomere.

### Telomere-associated protein genes exhibited similar conservation patterns among Panagrolaimidae species/isolates

We hypothesized that TTAGAC-containing unit clusters served as a component of partially stable telomere in TTAGGC-telomere species and that TBPs for TTAGGC telomere could have bound TTAGAC repeats and have been conserved in both TTAGGC-telomere and TTAGAC-telomere species. To support this hypothesis, we analyzed protein homologies in Panagrolaimidae with *B*. *xylophilus* as an outgroup species, in which proteins have been described to maintain or bind telomeres in *C*. *elegans* (Ahmed and Hodgkin 2000; Hofmann et al. 2002; Im and Lee 2003; Kim et al. 2003; Joeng et al. 2004; Im and Lee 2005; Meier et al. 2006; Boerckel et al. 2007; Raices et al. 2008; Meier et al. 2009; Ferreira et al. 2013; Shtessel et al. 2013; Dietz et al. 2021; Yamamoto et al. 2021). We identified 12 proteins that exhibited similar homology patterns between TTAGGC- and TTAGAC-telomere species/isolates (Supplemental Fig. S10). In contrast, four proteins (POT-2, DTN-2 (TEBP-2), HPR-9 and MRT-1) displayed inconsistent conservation patterns. POT-2 was conserved in only two of the three TTAGGC-telomere species/isolates, and DTN-2 (TEBP-2) remained in only one of the three TTAGAC-telomere species/isolates. HPR-9 was observed in only one TTAGAC-telomere isolate. MRT-1, required for telomerase activity in *C*. *elegans* (Meier et al. 2009), was not detected in *B*. *xylophilus* or in any of the three TTAGAC-telomere species/isolates but was present in all three TTAGGC-telomere species/isolates. However, MRT-1 functional domains were not conserved in the TTAGGC-telomere species/isolates.

## DISCUSSION

TRMs and their protein partners are well conserved and mostly co-evolving because they must be connected to maintain the DNA–protein complex, telomeres (Shakirov et al. 2009; Steinberg-Neifach and Lue 2015; Sepsiova et al. 2016; Červenák et al. 2019). Nevertheless, TRMs have evolved across several taxa (Fulnečková et al. 2013; Garavís et al. 2013; Peska and Garcia 2020; Červenák et al. 2021). In this study, we identified two novel TRMs in Nematoda and confirmed that TTAGAC, one of the two novel TRMs, composes telomeres in a subset of Panagrolaimidae isolates. We also hypothesized that TTAGAC evolved through TTAGAC-containing unit clusters and the robustness of the TTAGGC-suited TBPs to bind tandem arrays of TTAGAC repeats in a canonical TTAGGC-telomere ancestor.

The consistent pattern of TRM changes observed in other multicellular organisms supported our hypothesis. First, sequence changes in plant TRMs also exhibit key characteristics (Peska and Garcia 2020). In particular, the basal TRM in plants, TTTAGGG, (*Arabidopsis*-type) (Richards and Ausubel 1988) evolved into _TTAGGG (vertebrate-type) (Weiss and Scherthan 2002; Sýkorová et al. 2003; Sýkorová et al. 2006), <u>T</u>TTTAGGG (*Chlamydomonas*-type) (Fulnečková et al. 2012), TT<u>C</u>AGG_ (*Genlisea*-type) (Tran et al. 2015), <u>T</u>TT<u>C</u>AGG_ (*Genlisea*-type) (Tran et al. 2015), <u>T</u>TTTAGG_ (*Klebsormidium*-type) (Fulnečková et al. 2013) and <u>TTT</u>TTTAGGG (*Cestrum*-type) (Peška et al. 2015). Briefly, these variant TRMs exhibited alterations in the lengths of the T and G arms or changes in pyrimidine (T to C) nucleotides close to the middle A. Pyrimidine nucleotide conversion, rather than changes between pyrimidine and purine nucleotides and vice versa, has also been observed in insects in which TTAGG changed to T<u>C</u>AGG (Mravinac et al. 2011). Similarly, in the current study, TTAGGC changed to TTAGG<u>T</u> (pyrimidine conversion) and TTAG<u>A</u>C (purine conversion) in nematodes. Considering pyrimidines (T and C) have comparable base sizes and purines (G and A) are of similar sizes, we propose that conversions between pyrimidines or between purines do not sterically hinder TBP binding, allowing pre-existing TBPs to attach to the modified TRM. Furthermore, in yeasts and plants, TBPs are sufficiently robust to bind to various types of TRM despite several base pair changes (Rotková et al. 2004; Fajkus et al. 2005; Kramara et al. 2010; Visacka et al. 2012; Tomáška et al. 2019; Červenák et al. 2021).

It is still unclear whether TTAGGC-binding TBPs also bind to TTAGAC in TTAGGC-telomere Panagrolaimidae species/isolates. Traces of ALT action identified in our genome assemblies revealed that TTAGAC-containing units were used to repair telomere damage in TTAGGC-telomere isolates. Two TTAGGC-telomere isolates and one TTAGAC-telomere isolate examined harbored both clusters of consecutive TTAGGC- and TTAGAC-containing units.

Some clusters were attached directly or were adjacent to the telomeres. Specifically, the subtelomeric sequences of ptg000247l in LJ2406, a TTAGGC-telomere isolate, showed evidence of direct usage of a TTAGAC-containing unit for the repair of a damaged telomere. In the vicinity of a damaged and shortened telomere, there were homologous sequence blocks and a TTAGAC-containing unit cluster, which were probably replicated by BIR and MMBIR, and the TTAGAC-containing unit cluster was directly adjacent to its telomeric TTAGGC cluster. Thus, at least over a brief period of time, the TTAGAC-containing unit cluster was exposed at the end of the chromosome before the TTAGGC repeats were replenished.

Furthermore, the TTAGAC-containing unit cluster was not only duplicated but also elongated at the end because it was nearly twice as long as its initial cluster template in a separate subtelomeric region. This cluster might have been elongated via replication slippage based on its repetitive nature or via ALT mechanisms that can replicate TRM-containing units at the end of the chromosome. We could not determine the specific mechanisms involved in the elongation of the TTAGAC-containing unit cluster. If the elongation was achieved via ALT mechanisms, this suggests that the TTAGAC-containing unit cluster exhibits its own replication capability in the telomeric region to maintain the telomere.

If the TTAGAC-containing unit cluster acted as a component of the telomere, the TTAGGC-suited TBPs would bind to the telomere; otherwise, the end would be recognized as a DNA damage site. This binding robustness of TBPs is a plausible hypothesis in TRM evolution, and this robustness could be achieved via two potential ways: (1) the TTAGGC-suited TBPs also exhibited sufficient binding affinity for TTAGAC, such that TTAGAC-unit clusters could be utilized for repairing telomere damage. This binding affinity stabilized the TTAGAC-containing unit–TBP complex as a telomere. (2) TBPs exhibited a weak binding affinity to TTAGAC and formed a partially stable telomere complex, and the affinity strengthened through frequent use of TTAGAC-containing unit clusters to repair telomere damage and ensure fitness. In both cases, TTAGGC-suited TBPs could also stabilize TTAGAC telomeres, which thus facilitated the conversion from TTAGGC telomeres to TTAGAC telomeres.

Our results suggest a plausible evolutionary model for the TRM conversion in the family Panagrolaimidae. First, ancestral TTAGGC-telomere species may have utilized various ALT mechanisms to counteract telomere damage (Fig. 4). Replication of TTAGRC-containing unit clusters could represent one such ALT mechanism and would have been used to repair damaged telomeres. Because TBPs can fully or partially stabilize telomere complexes consisting of TTAGAC-containing unit clusters, the evolution of the TRM conversion to TTAGAC had a greater fitness than other TTAGGC variants. TRM would have converted to TTAGAC via mutation in the telomerase RNA component gene *terc*, leading to TTAGAC telomeres.

**Figure 4.**
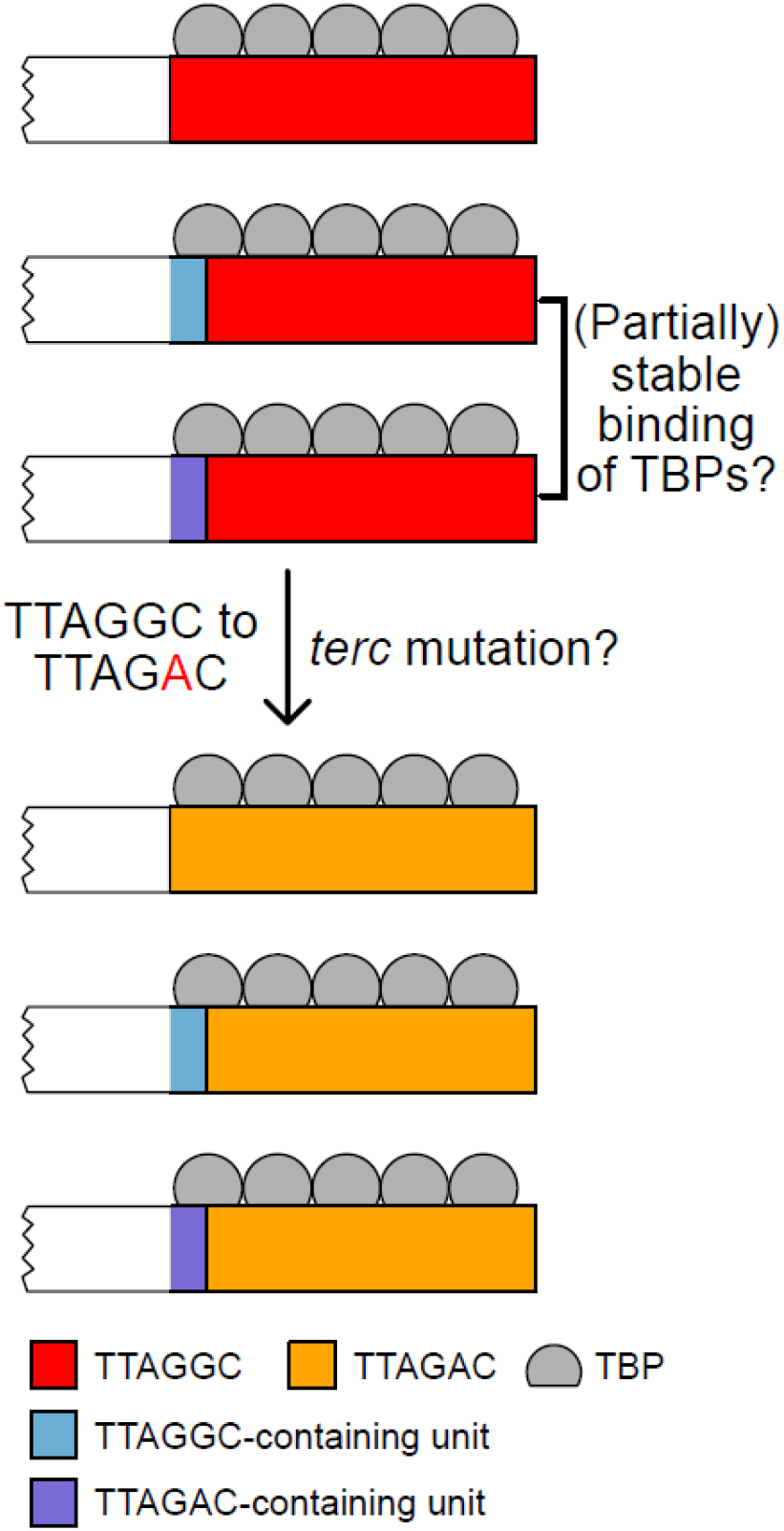
A model proposing TRM evolution in Panagrolaimidae through the TTAGAC-containing unit clusters and binding robustness of TBPs. TBPs with affinity for TTAGGC telomeres may bind to any of the TTAGGC telomere, TTAGGC-containing unit clusters, or TTAGAC-containing unit clusters, leading to a fully or partially stable telomere with TTAGRC-containing unit clusters. This binding robustness of TBPs may have facilitated the TRM conversion from TTAGGC to TTAGAC. Even following TRM changes by mutation of the telomerase RNA component gene (*terc*), TBPs can still bind to altered telomeric repeats, which contributed to the evolution of a novel TRM, TTAGAC, in the Panagrolaimidae family.

It is unclear whether TTAGGC-suited TBPs could bind to the novel telomeric sequence; however, if this were possible, the TTAGAC cluster–TBP complex would be partially stabilized, preventing the new telomere from being recognized as a DNA damage site. This hypothesis can be evaluated directly by inducing a mutation in the *terc* gene and then measuring the robustness of TBP in terms of sequence specificity. Unfortunately, we could not identify a *terc* gene for this species, as no *terc* gene has been identified even in *C*. *elegans*. Further studies are required to support our hypothesis. Nonetheless, we identified TTAGGC- and/or TTAGGT-containing unit clusters in putative subtelomeric regions of *Caenorhabditis uteleia*, which suggests that TRM-containing unit clusters are associated with telomere maintenance and that their usage in telomere maintenance contributes to TRM evolution.

Our results describe the independent evolution of two novel TRMs in Nematoda. We found that an ALT-mediated mechanism may have been used to repair ancestral telomere damage in Panagrolaimidae nematodes. Further, we hypothesized that the use of TTAGAC-containing units and the robustness of TPBs facilitated the evolution of the novel TRM. As the robustness of the TBPs may depend on the same-sized bases between canonical and novel TRMs, the evolutionary path we propose for Panagrolaimidae can also be applied to plants, insects or other nematodes where mutations occur only between pyrimidine nucleotides or between purine nucleotides. Although our study cannot explain all TRM changes in Nematoda, it provides new insight into telomere evolution through ALT and the robustness of TBPs.

## METHODS

### Worm sampling and culture

Nematodes were collected from rotten fruits in the Republic of Korea. They were grown in plates containing nematode growth medium seeded with *Escherichia coli* strain OP50 and contaminated by their natural microbes.

### DNA/RNA extraction and sequencing

Mixed-stage worms were lysed, and genomic DNA was extracted using the Gentra Puregene Cell and Tissue Kit (QIAGEN) to obtain WGS data for Panagrolaimidae isolates, LJ2281, LJ2284, LJ2285, LJ2400, LJ2402 and LJ2406. Macrogen (South Korea, https://www.macrogen.com/en/main) prepared DNA sequencing libraries using TruSeq Nano DNA and performed 151-bp paired-end DNA sequencing on the Illumina NovaSeq 6000 platform. One adult female worm was lysed, and its genomic DNA was amplified in the same 0.2-mL PCR tube using the REPLI-g single-cell WGA kit (QIAGEN) to obtain single-worm WGS data from LJ2050, LJ2051, LJ2060, LJ2070, LJ2072, LJ2401, LJ2411 and LJ2417. Paired-end DNA sequencing was performed using the Illumina NovaSeq 6000 platform by Theragen Bio (South Korea, https://www.theragenbio.com/en/).

Long-read DNA sequencing and short-read RNA sequencing of LJ2284, LJ2285, LJ2400 and LJ2406 were conducted as described by Kim et al. (Kim et al. 2019a) and Lee et al. (Lee et al. 2022a). In summary, genomic DNA was extracted from mixed-stage worms using phenol/chloroform/isoamyl alcohol (25:24:1) and sequenced using the HiFi mode of the PacBio Sequel II platform by Macrogen. RNA was extracted from mixed-stage worms using the TRIzol method. RNA sequencing libraries were prepared using TruSeq Nano DNA and sequenced by Macrogen using the Illumina NovaSeq 6000 platform with paired-end reads.

### Preparing publicly available WGS data

We included Nematoda species in our study, whose genome assembly data were available in the GenBank database and short-read sequencing data were available in the NCBI Sequence Read Archive (SRA) (RRID:SCR_004891). The following criteria were used to filter the short-read sequencing data: Instrument, *Illumina* (if HiSeq data were available, we used HiSeq; otherwise, MiSeq, Nextseq and Genome Analyser II data were selected); Source, *Genomic*; Layout, *PAIRED*; Number of Bases, *>5 Gb*; Spot number, *>5 million*; Read length, *>70 bp*. Nematode clades were classified according to the phylogenetic tree (Smythe et al. 2019). Any genus that was not included in the phylogenetic tree was excluded. WGS data for *Enoplus brevis,* which lacked genome assembly information in the GenBank database, and WGS data for *Panagrellus redivivus* (Srinivasan et al. 2013) that was supplied by Dr. P. W. Sternberg were added to the filtered datasets. Detailed accession information is available in Supplemental Tables S1 and S2.

### Identifying TRMs using short-read WGS data

To normalize the public WGS data, we trimmed all reads to 60 bp and used only 5 million sub-sampled reads from the R1 files using Seqtk (version 1.3-r106; *seqtk sample -s 11 - 5000000*). For each dataset, 23-mers were counted using Kounta (version 0.2.3; *kounta -- kmer 23 --out*). Only tandemly repeated sequences with unit lengths of 5–7 bp and counts >99 were used to identify putative TRMs that were most frequent and most similar to the canonical TTAGGC sequence. Methods used to further validate the novel putative TRMs are described in Supplemental Methods. We averaged the counts of the 23-mers that contained the corresponding TRM concatemers to compare the number of TRMs across species. For the Panagrolaimidae isolates, we used 20 million sub-sampled reads and all 23-mers, as telomeric repeats were not evenly amplified during whole-genome amplification.

### Genome assembly

HiFi reads were *de novo* assembled using Hifiasm (Cheng et al. 2021) (version 0.13-r308; *hifiasm -l0*). Contigs that were potentially contaminated with bacterial sequences were removed as described by Kim et al. (Kim et al. 2019a). Scaffolding and visualization based on the Hi-C data are described in Supplemental Methods. The completeness of the genome assembly was evaluated using BUSCO (Simão et al. 2015) (version 4.0.6; *busco -m genome -l nematoda_odb10*) with the nematoda dataset in the OrthoDB release 10. Isolate-specific repetitive sequences were identified using BuildDatabase (Flynn et al. 2020) (version 2.0.1; default options) and RepeatModeler (Flynn et al. 2020) (version 2.0.1; *RepeatModeler - database -LTRStruct*). Isolate-specific and known metazoan-repetitive sequences were masked using RepeatMasker (Chen 2004) (version 4.1.0; *RepeatMasker -lib -s* for species-specific repeats and *RepeatMasker -species metazoa -s* for metazoan repeats). We mapped the RNA-seq reads to the repeat-masked genome using HISAT2 (Kim et al. 2019b) (version 2.2.1; *hisat2-build* for genome indexing and *hisat2* with default options to map the RNA-seq reads to the corresponding genome). RNA-mapping data was used to annotate genes using BRAKER (Stanke et al. 2006; Stanke et al. 2008; Li et al. 2009; Barnett et al. 2011; Lomsadze et al. 2014; Buchfink et al. 2015; Hoff et al. 2016; Hoff et al. 2019; Brůna et al. 2021) (version 2.1.5; *braker.pl --genome -bam --softmasking*).

### Preparation of 18S ribosomal DNA sequences to generate the Panagrolaimidae phylogenetic tree

For *Panagrolaimus* sp. PS1159, *Panagrolaimus davidi*, *Panagrellus redivivus* and *Bursaphelenchus xylophilus*, we used publicly available 18S rDNA sequences (GenBank accessions: U81579.1, AJ567385.1, AF083007.1 and KJ636306.1, respectively). For the HiFi-based *de novo* genome assemblies, we extracted 18S rDNA sequences by searching known nematode 18S rDNA PCR primers (nSSU_F_04: 5′-GCTTGTCTCAAAGATTAAGCC-3′ (Blaxter et al. 1998) and nSSU_R_82: 5′-TGATCCTTCTGCAGGTTCACCTAC-3′ (Medlin et al. 1988)) in genome assemblies using BLAST+ (Camacho et al. 2009) (version 2.7.1; *makeblastdb - input_type fasta -dbtype nucl* and *blastn -task blastn-short -outfmt 6*).

For the other TTAGAC-TRM isolates used in this study, we mapped their short-read DNA sequencing data to the LJ2285 genome assembly using BWA-MEM (version 0.7.17; *bwa mem*, default option), and the 18S rDNA sequence variation of each isolate were identified using BCFtools (Li 2011a) (version 1.13; *bcftools mpileup -Ou -f | bcftools call -Ou -mv | bcftools norm -f -Oz -o*). The indexed output VCF files were obtained using Tabix (Li 2011b) (version 1.13; default option), and the LJ2285 18S rDNA sequence was replaced with their variants of each isolate using SAMtools (Li et al. 2009) (version 1.13) and BCFtools (*samtools faidx -r | bcftools consensus -o*). For the other TTAGGC-TRM isolates, we used the LJ2400 genome assembly as a reference and repeated the procedure described above. All 18S rDNA sequences of Panagrolaimidae obtained in this study are listed in Supplemental Table S6. We generated an alignment file using all 18S rDNA sequences as input for Clustal Omega (https://www.ebi.ac.uk/Tools/msa/clustalo/, RRID:SCR_001591, (Sievers and Higgins 2021)) (Sequence Type: DNA; Output Alignment Format: PHYLIP). We used this alignment as an input to RAxML (Stamatakis 2014) (version 8.2.12) using raxmlGUI 2.0 (Edler et al. 2021) (version 2.0.10; options: GTRGAMMA and ML+rapid bootstrap with 1000 replications) to infer the phylogenetic relationships, which were visualized using Dendroscope (Huson and Scornavacca 2012) (version 3.8.4).

### Telomere and subtelomere structure in the genome assemblies

We selected candidate telomere-containing contigs using two tandemly repeated copies of TTAGRC that appeared ≥3 times in the 600-bp region at each end of the contig. We manually validated these telomeric regions using the following criteria: TRM cluster length ≥500 bp (the longest ITS length in *C*. *elegans*) (Yoshimura et al. 2019) or TRM cluster still located at the end after scaffolding (only for LJ2284 and LJ2406). We analyzed whether each telomere-containing contig/scaffold had clusters with ≥6 copies of TRMs or whether TTAGRC-containing units repeated tandemly in its subtelomeric region (up to 200 kb from the end of the contig/scaffold). For LJ2285, we considered sequences consisting of 5–9 copies of different units that contained TTAGRC and similar sequences as unit clusters because LJ2285 did not contain clusters composed of the same units. Subsequently, we investigated whether the unit cluster (containing ≥6 copies of units, the same TTAGRC and identity ≥85% of units) existed outside the subtelomeric region of telomere-containing contigs/scaffolds using BLAST+ (Camacho et al. 2009) (version 2.12.0; *makeblastdb - input_type fasta -dbtype nucl*; *blastn -task blastn-short -outfmt 6* for sequences <30 bp and *blastn -task megablast -outfmt* for sequences ≥30 bp). Unit sequences of LJ2284, LJ2400 and LJ2406 and unit cluster sequences in LJ2285 were denoted starting with TTAGRC (Supplemental Table S12). The break-induced replication (BIR) traces were analyzed by searching for homologous sequences between ITS, TTAGRC-containing unit cluster, or telomeric repeats against their corresponding genome assembly using BLAST+ (Camacho et al. 2009) (version 2.7.1; *makeblastdb -input_type fasta -dbtype nucl and blastn -task megablast -outfmt 6*).

### Conservation analysis of telomere-associated proteins

Protein FASTA sequences were downloaded from WormBase for *C*. *elegans* (Release WS281) and WormBase ParaSite (Howe et al. 2017) for *Panagrolaimus* sp. PS1159, *Panagrolaimus davidi*, *Panagrellus redivivus* and *B. xylophilus* (Release WBPS15). Protein sequences for *Panagrolaimus* sp. PS1579 and *Panagrolaimus* sp. ES5 were not publicly available. For the four Panagrolaimidae isolates (LJ2284, LJ2285, LJ2400 and LJ2406), we used protein FASTA sequences obtained through gene annotation using BRAKER. For the non-*C*. *elegans* nematodes, we searched for protein sequences that were conserved in *C*. *elegans* using DIAMOND (Buchfink et al. 2021) (version 2.0.11; *diamond blastp -d -q -o --threads 20 --very-sensitive* and *diamond blastp -d -q -o --threads 20 --ultra-sensitive*) to characterize telomere-associated protein sequences that included TRT-1, POT-1, POT-2, POT-3, MRT-1, MRT-2, DTN-1 (TEBP-1), DTN-2 (TEBP-2), HPR-9, HPR-17, HUS-1, SUN-1, CEH-37, HMG-5, HRPA-1 and PLP-1. We filtered the nematode’s highest bit score protein sequence for each *C*. *elegans* telomere-associated protein and identified the conserved domains using NCBI CD-search (Marchler-Bauer and Bryant 2004; Lu et al. 2020) (CDD database, version 3.19).

## DATA ACCESS

Raw sequencing data and genome assemblies have been deposited in the NCBI BioProject database (https://www.ncbi.nlm.nih.gov/bioproject) under the accession number PRJNA845886 and in the Korean Nucleotide Archive (KoNA, https://kobic.re.kr/kona) under the BioProject accession number PRJKA220348 (sequencing data only).

## ACKNOWLEDGEMENTS

This study was supported by Samsung Science and Technology Foundation [SSTF-BA1501-52]; and National Research Foundation of Korea [2019R1A6A1A10073437 to J.K.]. J. Lim was also supported by a scholarship for basic research, Seoul National University. We thank Dr. P. W. Sternberg for providing valuable sequencing data of *Panagrellus redivivus*. We also thank Namhee Kim and Sungyeol Ahn for providing accession to their farms to collect rotten fruits and Dr. D. S. Lim for collaboration in collection of nematode species. We appreciate Dr. S. Sung for providing the suggestions on generating Hi-C library, and Dr. C. Kim for the help in Hi-C data processing.

## AUTHOR CONTRIBUTIONS

Jiseon Lim: Conceptualization, Methodology, Formal Analysis, Investigation, Writing-Original Draft, Writing-Review & Editing. Wonjoo Kim: Investigation. Jun Kim: Conceptualization, Methodology, Investigation, Writing-Original Draft, Writing-Review & Editing. Junho Lee: Conceptualization, Writing-Review & Editing, Funding Acquisition, Supervision.

